# Rolling stones gather moss: Movement and longevity of moss balls on an Alaskan glacier

**DOI:** 10.1101/687665

**Authors:** Scott Hotaling, Timothy C. Bartholomaus, Sophie L. Gilbert

## Abstract

Glaciers support diverse ecosystems that are largely comprised of microbial life. However, at larger, macroscopic scales, glacier moss balls (sometimes called “glacier mice”) can develop from impurities on ice surfaces and represent a relatively rare biological phenomenon. These ovoid-shaped conglomerations of dirt and moss are only found on some glacier surfaces and provide key habitats for invertebrate colonization. Yet, despite their development and presence being widely reported, no targeted studies of their movement and persistence across years have been conducted. This knowledge gap is particularly important when considering the degree to which glacier moss balls may represent viable, long-term biotic habitats on glaciers, perhaps complete with their own ecological succession dynamics. Here, we describe the movement and persistence of glacier moss balls on the Root Glacier in southcentral Alaska, USA. We show that glacier moss balls move an average of 2.5 cm per day in herd-like fashion, and their movements are positively correlated with glacier ablation. Surprisingly, the dominant moss ball movement direction does not align with the prevailing wind or downslope directions, nor with any dominant direction of solar radiation. After attaining a mature size, glacier moss balls persist for many years, likely in excess of 6 years. Finally, we observed moss ball formation on the Root Glacier to occur within a narrow, low albedo stripe downwind of a nunatuk, a potential key source of moss spores and/or fine-grained sediment that interact to promote their formation.

## Introduction

Glaciers have long been overlooked as important components of global biodiversity (Stibal et al. 2020), but it is now clear that they host thriving, multi-trophic ecosystems (Anesio and Laybourn-Parry 2012), supporting taxa from microbes to vertebrates (Rosvold 2016; Dial et al. 2016; Hotaling et al. 2017a; Hotaling et al. 2019). Most biological activity on glaciers occurs within surface ice where microorganisms take advantage of nutrients that are either wind-delivered or generated *in situ* (Hotaling et al. 2017a). In addition to a nutrient input, impurities on the glacier surface can drive the development of at least two potential “hotspots” of biological diversity on glaciers: well-studied cryoconite holes (depressions in the ice surface caused by local melt, Anesio et al. 2017) and glacier moss balls (ovular conglomerations of moss and sediment that move on the glacier surface, Coulson and Midgley 2012).

Often a small piece of rock or other impurity sets in motion the formation of a glacier moss ball [also referred to as “jokla-mys” (Eythórsson 1951), “glacier mice” (e.g., Coulson and Midgley 2012), or “moss cushions” (e.g., Porter et al. 2008)]. On a local scale, glacier moss balls are typically distributed with some degree of local clustering (e.g., ∼1 glacier moss ball/m^2^; Fig. 1). While immobile moss aggregations have been observed on glaciers elsewhere (e.g., East Africa, Uetake et al. 2014), true glacier moss balls appear to be particularly rare, having only been described on a few geographically disparate glaciers in Alaska (Shacklette 1966; Heusser 1972), Iceland (Eythórsson 1951), Svalbard (Belkina and Vilnet 2015), and South America (Perez 1991). Many different moss species have been found in glacier moss balls (Shacklette 1966; Heusser 1972; Perez 1991; Porter et al. 2008), suggesting that they are not dependent on specific taxa, but instead their development is driven by the interaction of suitable biotic (e.g., availability of moss spores) and abiotic (e.g., growth substrate) factors. However, the specific steps and timeline of glacier moss ball genesis remains unclear.

**Fig. 1.**
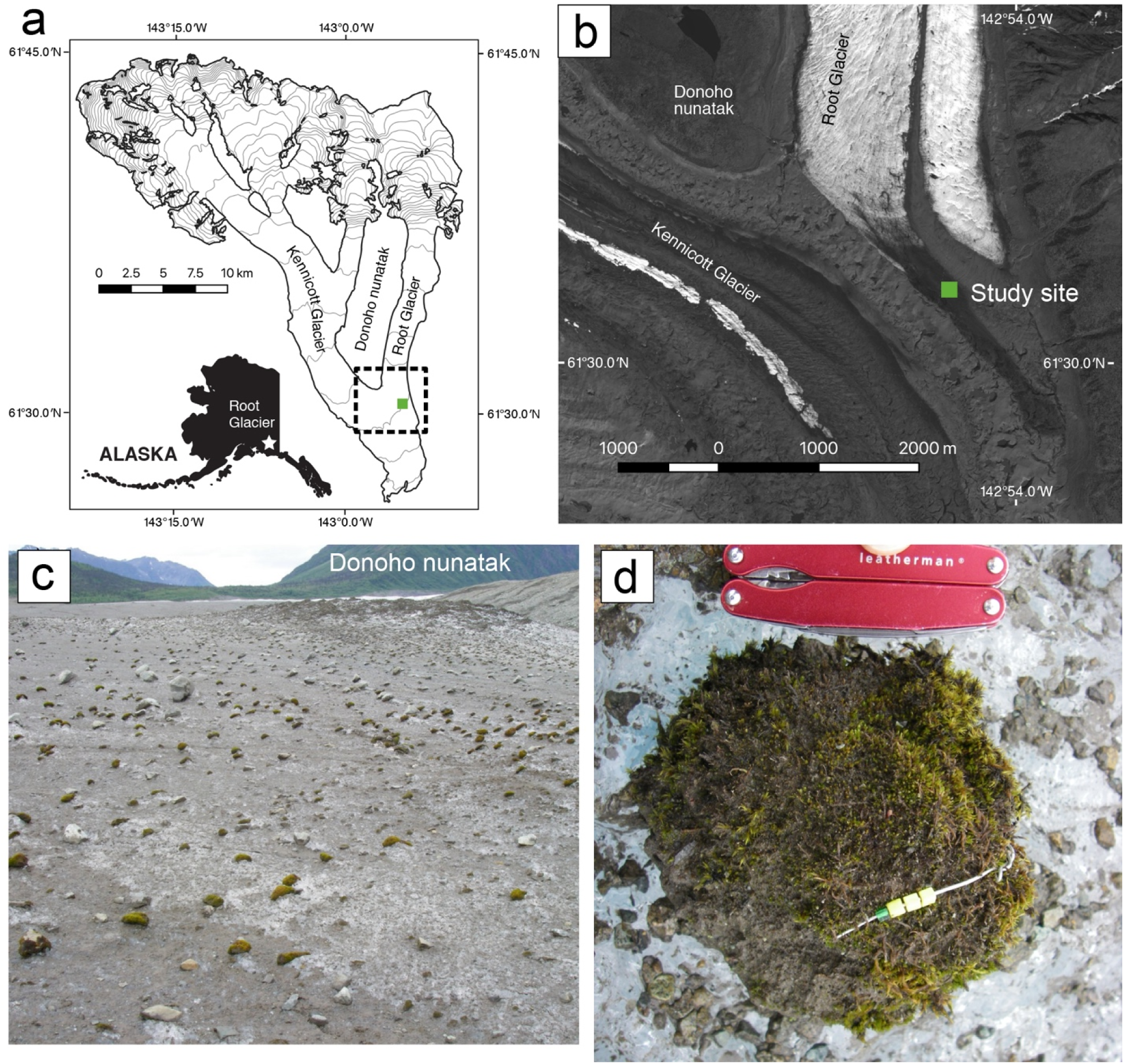
a) Our study site (solid green square) on the Root Glacier in southcentral Alaska, USA, within Wrangell-St. Elias National Park. Contour lines are spaced every 100 m in elevation. The dashed square represents the field of view shown in panel (b). The inset map shows the location of the Root Glacier (white star) within Alaska. b) Satellite image of the study site (green square) showing the confluence of the Root and Kennicott Glaciers with the Donoho nunatak to the northwest. The image was recorded on 19 June 2013. c) A landscape view looking northwest of the study site dotted with glacier moss balls. d) A close-up view of a glacier moss ball with the type of bracelet tag used in this study.

An intriguing aspect of glacier moss balls, and one that is almost certainly partially responsible for their “glacier mice” namesake, is their movement. It has been posited that moss balls move by inducing the formation of an ice pedestal, then rolling or sliding off of it (Porter et al. 2008). Under this process, moss balls first reduce local albedo by shielding the ice beneath them from sunlight and locally reducing the ablation rate. As the surrounding ice melts, the glacier moss ball is left on an elevated pedestal. Eventually, a threshold is reached where the moss ball falls from its pedestal and the process begins anew, potentially including a “flip” of the moss ball that exposes what was previously their underside (Porter et al. 2008). Though the speed and direction of moss ball movement has not been measured, though it has been suggested that their movements generally track the downslope direction of their local habitat (Porter et al. 2008).

Where they occur, glacier moss balls contribute to glacier biodiversity by offering a thermally buffered, island-like habitat on the glacier surface that hosts an array of invertebrates (Coulson and Midgley 2012). On Icelandic glaciers, moss balls contain invertebrate communities dominated by springtails (Collembola), tardigrades (Tardigrada), and nematodes (Nematoda; Coulson and Midgley 2012). While many potential food resources are available on glaciers (Hotaling et al. 2017a, 2020), these are typically only exploited by invertebrates on the margins (e.g., springtails, spiders, grylloblattids), likely because suitable on-glacier habitat is lacking (Mann et al. 1980). Glacier moss balls may therefore provide key habitable islands on the glacier that facilitate wider resource exploitation versus glaciers without moss balls (Coulson and Midgley 2012). It is also possible that glacier moss balls, which have not been shown to be inhabited by larger predatory insects (e.g., grylloblattids) may provide prey refuge that are sufficiently removed from the typical foraging areas of their predators. Either way, it is clear that glacier moss balls represent important habitat for glacier-associated fauna yet basic aspects of their ecology (e.g., longevity and movement) are unknown.

In this study, we took an integrated behavioral ecology and geophysical approach to the study of glacier moss balls to answer three basic questions about their life history: (1) How long do mature glacier moss balls persist on the landscape? (2) How quickly do they move and is their movement idiosyncratic or herd-like? (3) Are the movements of glacier moss balls linked to the ablation of the glacier itself? Answers to these questions have implications for invertebrate fauna in glaciated ecosystems, nutrient cycling (both directly via moss ball decomposition and indirectly as supporting habitat for biotic communities), and feedback between glacier moss balls and local ablation rates. Beyond biotic interactions and ecosystem dynamics, glaciers are rapidly receding worldwide (Gardner et al. 2013; Larsen et al. 2015; Roe et al. 2017) and their diminished extents will almost certainly affect the persistence of glacier moss balls on local and global scales. Thus, it is important to better understand these unique micro-ecosystems before their habitats are lost.

## Materials and methods

### Study area

We conducted fieldwork over four years (July, 2009 - July, 2012) on the lowest portion of the Root Glacier, a major tributary to the Kennicott Glacier, in the Wrangell Mountains in Wrangell-St. Elias National Park, Alaska, USA (Fig. 1a). Our study area (61.5076° N, 142.9172° W, ∼700 m elevation) spanned a ∼15 x ∼40 m (600 m^2^) area of glacier ice selected for its especially high concentration of moss balls. The site has a gentle slope, dipping 3° east-northeast (N75°E) and is found between two medial moraines (Fig. 1b), each ∼100 m away. Glacier surface speeds here are slow, typically 0.05 to 0.15 m d^-1^ during summer (Armstrong et al. 2016). Several, narrow (<1 cm wide) and stagnant crevasses (manifesting as closed, linear, surface depressions) cross our study area, but did not significantly disrupt the otherwise consistent slope of the site. Moss ball concentrations decrease both up- and down-glacier and are absent from the coarse-grained (> 5 cm) rock that covers the adjacent medial moraines.

We estimated the proportion of fine-grained sediment cover on the ice within our study area by applying image processing techniques in the Python package scikit-image (Van der Walt et al. 2014) to two vertical photographs taken of representative ice surfaces. Pixel brightness contrasts between ice and sediment are most distinct within the blue band of the red-green-blue images, so we differentiated between sediment (dark pixels) and ice (bright pixels) by binarizing the blue band with Otsu’s thresholding method. We then performed a morphological opening to diminish the influence of light-colored sediment grains set within the otherwise dark sediment cover. Finally, we quantified the areal sediment cover as being approximately equal to the number of dark colored pixels relative to the total number of pixels in the binarized images.

### Mark-recapture

During the summer of 2009, we tagged 30 glacier moss balls with a bracelet identifier (Fig. 1d). We focused our efforts on “mature” moss balls that had reached at least ∼10 cm in length on their longest axis and were ovoid with no obvious morphological irregularities. Each bracelet consisted of a unique combination of colored glass beads (∼2-3 mm in diameter) threaded on aluminum wire. Bracelets were threaded through the moss ball center and pulled snug so as to not protrude beyond the moss ball’s exterior and interfere with movement. We returned eight times during the 2009 season to re-survey moss balls and record their movements. We followed up our initial surveys with annual visits from 2010-2012. During each survey, we visually inspected in and around the core study area multiple times in an effort to recapture moss balls. As part of this process, we visually inspected each moss ball in the area for any sign of a bracelet tag. After inspection, we replaced each moss ball in the exact location and orientation as it was found.

### Moss ball movement and glacier ablation

We assessed moss ball movement over 54 days in 2009. As benchmarks for their movement, we installed three ∼1.3 cm PVC tubes into the glacier. Each stake was drilled ∼60 cm into the glacier. Stakes were installed in a triangle that spanned the study area and served two purposes. First, the stakes provided a reference against which the location of each moss ball was measured. Second, they allowed us to measure glacier ablation (i.e., the distance the ice surface moves vertically down) over the same study period so we could test for links between moss ball movement and the rate of glacier ablation.

To track glacier moss ball movement, during each site visit, we measured the distance between re-identified moss balls and each reference stake with a flexible, fiberglass measuring tape, pulled taught between the moss ball center and reference stake. Next, for each moss ball, we used trilateration to calculate three independent positions within our field site—one for each of the three pairs of reference stakes. We assigned the location of a surveyed moss ball to the mean of these three relative positions and constructed a location covariance matrix for each measurement, to assign uncertainties to surveyed locations. After diagonalizing the covariance matrix, we identified the size (eigenvalues) and orientation (eigenvectors) of an uncertainty ellipse around each mean location. Major and minor axes of the uncertainty ellipse were defined as twice the square root of the eigenvalue lengths, such that each error ellipse represented a 2σ error window. Thus, assuming independent, normal errors, we are 95% confident that the true location of each moss ball fell within its error ellipse. The size of each error ellipse thus accounts for potential errors including failure to pull the tape completely tight in the face of katabatic winds or long measurement distances, or inconsistent identification of moss ball centers. While we used stakes for most of the measurement period, we were forced to switch to washers (∼5 cm in diameter) laid flat on the ice surface later in the season, during a period when we were unable to drill the benchmark stakes sufficiently deep to avoid melting out between visits. Before transitioning from benchmark stakes to washers, we tested the stability of the washers to ensure that they did not slide over the ice surface. Over a 5-day period in early August, we did not detect significant washer movement (outside of 2σ uncertainty). Only the final measurements (11 August 2009) and calculations were made relative to the washers. From moss ball position data, we calculated mean speeds and azimuths (travel directions) between position measurements for each moss ball. Moss ball velocities are reported relative to a reference frame that travels with the ice surface, into which the reference stakes were drilled and onto which washers were placed. Velocities are therefore unaffected by bulk glacier motion. To quantify glacier ablation, the height of each stake above the local ice surface was re-measured during each visit and periodically re-drilled into the ice as necessary. Ablation reported in this study is the mean ice surface lowering rate calculated for each of the three stakes. As an assessment of ablation uncertainty, we also calculated the maximum deviation of any single stake’s ablation rate from the overall mean.

We assessed the potential for East/West asymmetry in the direction of incoming solar radiation as a control on the direction of moss ball movement using a time series of solar radiation from a Remote Automatic Weather Station (RAWS) located 15 km up-glacier from our study site and approximately 500 m higher in elevation. The RAWS site, at Gates Glacier (https://wrcc.dri.edu/cgi-bin/rawMAIN.pl?akAGAT), is located on a ridge above the Kennicott Glacier and records incoming solar radiation and other meteorological variables every hour. To evaluate the relative levels of solar energy arriving at our field site before and after solar noon, we integrated each afternoon’s solar radiation and subtracted each morning’s integrated solar radiation from it, thus arriving at a daily metric of the morning/afternoon solar energy asymmetry. Values near 0 indicated equal amounts of energy arriving during mornings and afternoons, positive values indicated more solar energy during the afternoons than mornings, and negative values revealed more incident energy during the mornings.

### Persistence

We sought to understand how long mature glacier moss balls persist on the landscape, particularly across years. We hypothesized that mature moss ball longevity might vary due to differences in environmental conditions (e.g., precipitation, freeze-thaw cycles) or random chance (e.g., a crevasse opening within a key area). Furthermore, we wanted to know not only how likely we are to detect glacier moss balls, given that they had persisted within the study area, but also if our detection probability varies among years. To do this, we fit capture-recapture models of annual survival to each glacier moss ball included in the study. Because moss balls were individually marked but were not equipped with radio-transmitters or other devices which would allow us to know their ultimate fates, we applied Cormack-Jolly-Seber (CJS; Lebreton et al. 1992) survival models. These CJS models develop a “capture history” of each moss ball to estimate apparent survival (i.e., the probability that an individual is in the population at time *i* and still in the population at time *i*+1) and probability of detection if they persisted within our study area. Survival estimates from CJS models only represent apparent survival because emigration cannot be estimated from survival data with unknown fates (i.e., we did not know if a tagged moss ball had disaggregated, lost its identifying bracelet, or was no longer in the study area). Therefore, our estimates of apparent survival are likely to underestimate true survival (e.g., a moss ball might have lost its bracelet or moved out of the study site). In addition, CJS models also account for imperfect detection. In our case, if a moss ball persisted within our study area but was overlooked.

Using our individual moss ball annual detection data (1 = detected, 0 = not detected), we fit four competing CJS survival models, including the null model [no effect of year on apparent survival (□) or detection probability (*p*); Model 1)], an effect of year on □ (Model 2), an effect of year on *p* (Model 3), or an effect of year on both □ and *p* (Model 4). We then selected the model(s) best supported by our data using Akaike’s information criterion (AIC; Akaike 1998), adjusted for small sample size (AICc; Hurvich and Tsai 1989). Our model selection approach was based on model likelihoods and models were penalized for extra parameters to favor parsimony.

Finally, we calculated the average life expectancy of a mature glacier moss ball. To do this, we used annual survival rates based on life-table analysis (Deevey Jr 1947; Millar and Zammuto 1983), in which average life expectancy was calculated as -1/ln(Annual Survival Rate). Because this estimation of life expectancy is quite sensitive to annual survival rate, we calculated it for both the lowest annual survival rate and the mean annual survival rate. Thus, the true average life expectancy might be substantially greater than the conservative values estimated here. This framework for estimating average life expectancy does not account for variable mortality rates when glacier moss balls are first forming or nearing the end of their lifespans.

## Results

### Study area

Our study area was located on a “bare ice” glacier surface, between two medial moraines covered by coarse-grained, angular, rock debris. However, two types of sediment distinguish the study area surface from what would be considered clean, pure, water ice. First, glacier moss balls were found amidst gravel and small boulders (< 30 cm diameter), spaced every ∼1 m. Second, the ice surface has an unusually pervasive, fine-grained sediment cover, ∼1-3 mm thick, which partially blankets the otherwise bare ice. Image processing indicated that this fine sediment covers approximately 70% of the study area surface. This low albedo sediment cover is visible in all inspected satellite imagery of the site and first appears at lower concentrations emerging from cleaner ice ∼1 km northwest of the study site (Fig. 1b). Down-glacier of the study site, the low albedo region extends ∼1.7 km as a ∼300-m-wide, rounded finger that spans adjacent medial moraines, in a manner consistent with wind-deposited dust, draping over underlying geomorphic features. Therefore, we interpreted the southeast (135°) trend direction of this low albedo finger to be the prevailing, down-glacier, katabatic wind direction. During the 26 days of glacier ablation measurements, the ice surface lowered by 1.91 m due to melt and sublimation. Ablation rates ranged from 5.8-9.6 cm per day (cm d^-1^) between measurement times and averaged 7.3 cm d^-1^.

### Movement

Glacier moss ball movements varied systematically over the study period, with increases and decreases that coincided with changes in direction (Figs. 2-3). Median moss ball speed was 2.5 cm d^-1^, but their rates varied widely throughout the season. The median speed started at 1.8 cm d^-1^ in late June, increased to 4.0 cm d^-1^ at the start of July, then slowed to 2.0 cm d^-1^ during late July/early August. The maximum observed speed for any glacier moss ball was 7.8 cm d^-1^ during the 5-day period from July 9-14 (excluding two outlier speeds that were more than 8 interquartile ranges greater than the median, 14.2 and 21.0 cm d^-1^, and which were based upon particularly uncertain moss ball positions). The interquartile range of moss ball speeds was approximately 50% of the median speed; thus, these observed increases and decreases in speed reflect changes in the entire population of moss balls.

**Fig. 2.**
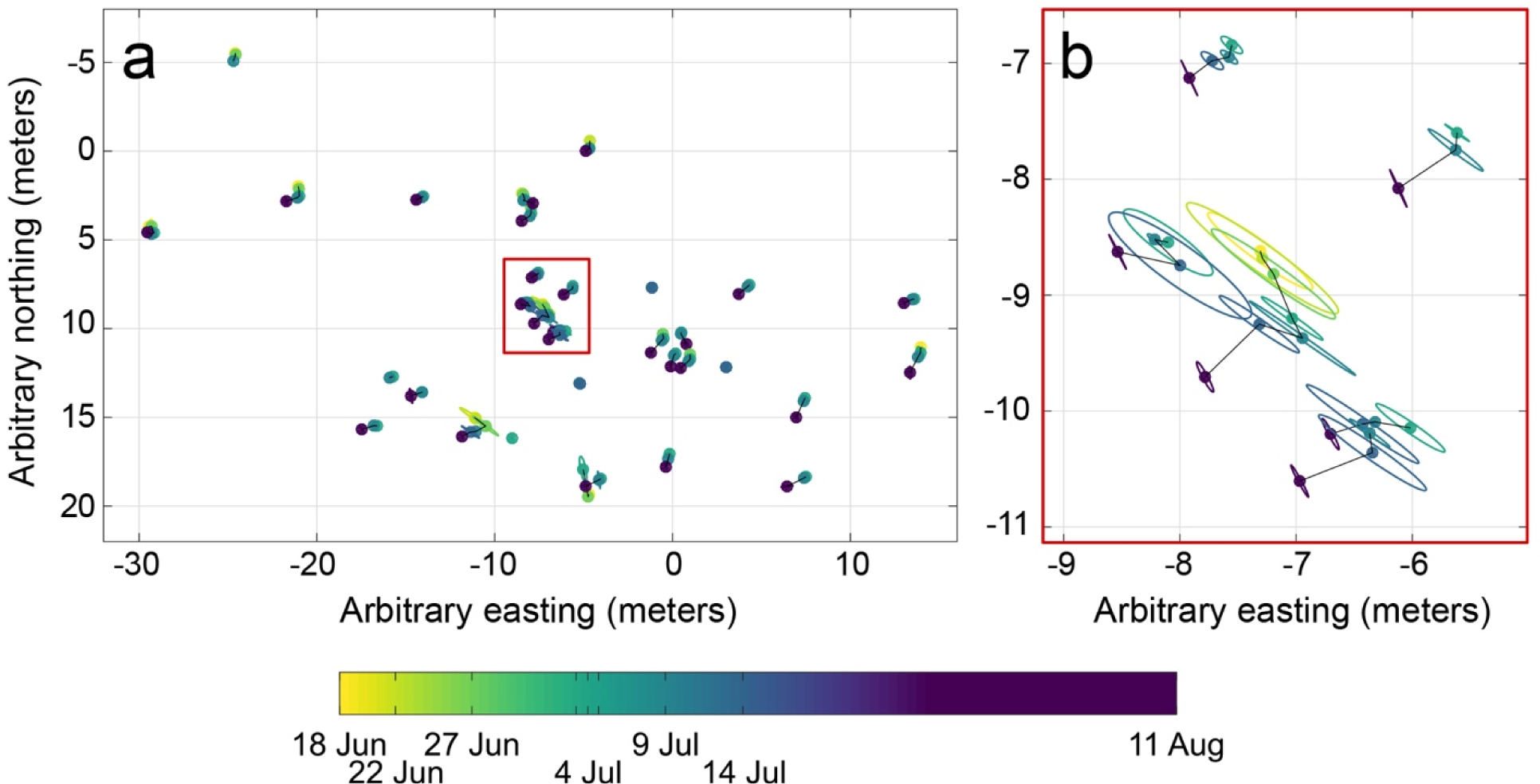
(A) Locations of surveyed glacier moss balls throughout the survey period. Most likely locations of each moss ball are shown with small filled circles relative to an arbitrary, local grid system. Ellipses surrounding each moss ball indicate 2σ uncertainty (i.e., 95% confidence) of their location. Thin black lines connect consecutive surveyed locations for individual moss balls. The red rectangle identifies the location of the large-scale view in panel (B). (B) A zoomed in view of movement patterns for six glacier moss balls (red square in A), showing their similar azimuths.

**Fig. 3.**
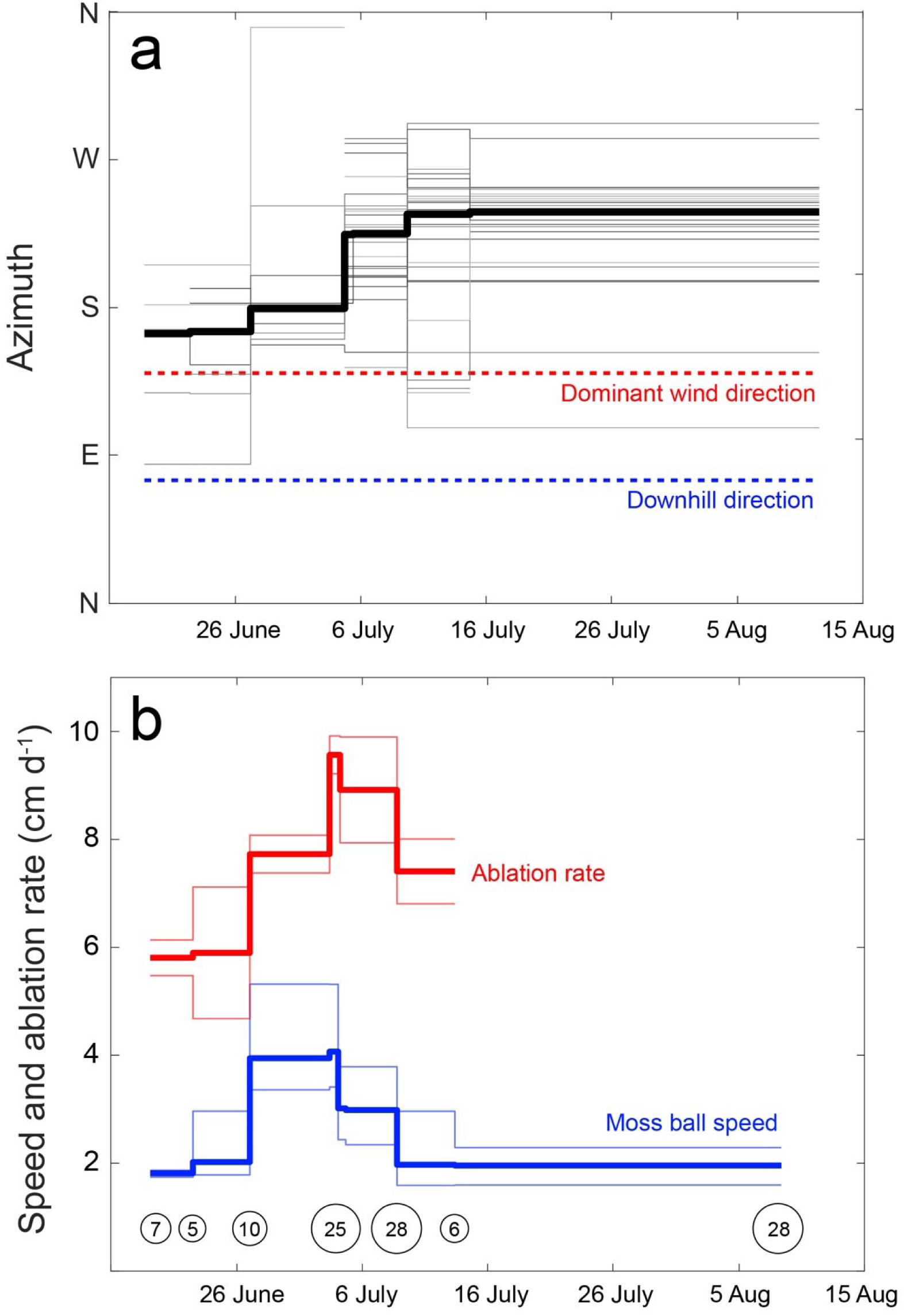
(A) A comparison of glacier moss ball movements versus the dominant wind (dashed red line) and downslope (dashed blue line) directions. Direction of each moss ball’s motion between measurement times is shown with thin gray lines, while the bold black line indicates the median direction of all glacier moss ball movements. (B) Glacier moss ball movement versus ablation rate. Median ablation rate is indicated with a bold red line, while the mean +/- the maximum absolute deviation from the mean are shown with thin red lines. The median speed of glacier moss balls is shown with the bold blue line, while the 25th and 75th percentile speeds are shown with thin blue lines. Numbers in circles along the bottom of the plot represent the number of moss balls surveyed at each timepoint (single measurements not indicated).

The direction of glacier moss ball movements was not random. Rather, glacier moss balls underwent clear changes in their direction of motion (i.e., azimuth) throughout the summer season (Fig. 3a). While individual moss balls moved in many directions, when viewed in aggregate, azimuths of the population clearly clustered over time. Early in the season, median moss ball motion was south-southeast (165°) but over the ensuing weeks azimuths progressively increased, such that at the end of the measurement period the median azimuth was west-southwest (240°; Fig. 3a).

Considering speeds and azimuths together, we see the moss ball population initially moving at 2 cm d^-1^ to the south for 9 days, then the group nearly doubles its speed to 4 cm d^-1^ while deviating slightly to the right (towards the west). After a week at these maximum speeds, speeds drop by 25% to 3 cm d^-1^ while also deviating 45 degrees further towards the west for five days. During the next 5-day measurement period, speeds drop further, back to 2 cm d-1 while the azimuths turn another 10-15 degrees further west. Over the final 28-day measurement period, the azimuths remain stable, while speeds continued to fall. This decrease in speed is apparent in the decline of the upper quartile of speeds, despite our not making sufficient new measurements to influence the median speed.

Our fine-scale movement and ablation data allowed us to compare glacier moss ball speeds and azimuths with potential drivers of their motion. We find that more rapid moss ball speeds are associated with more rapid ablation; an ordinary least squares model between ablation rate and speed indicates that, on average, for every 1 cm of surface ablation, the glacier moss balls move horizontally 0.34 cm (Fig. 3b). However, the relationship between ablation rate and speed is relatively weak (R^2^ = 0.40). It should also be noted that during the course of our study, participants in a program hosted by the Wrangell Mountains Center, McCarthy, Alaska, visually confirmed the posited primary movement method described by Porter et al. (2008), when a glacier moss ball was observed rolling off its elevated pedestal and inverting in the process.

The directions of moss ball motion, however, are more puzzling. The southern and western directions of moss ball movement are clearly distinct from both the prevailing, katabatic wind direction as inferred from the dust plume (towards the southeast) or the downhill direction of the gently sloping ice surface (towards the east-northeast; Fig. 3a). The herd-like change in travel direction, from an initially southerly direction to a southwesterly direction late during our measurement period, could potentially be explained by a shift in the dominant direction of incoming solar radiation. If, during the latter portion of July and August, 2009, the afternoons were sunnier than the mornings, then we would expect faster ice surface lowering on the southwest side of moss balls than on their northeast sides, and the moss balls would be more likely to roll off their ice pedestals towards the southwest, as observed. However, our analysis of solar radiation measurements revealed no such asymmetry (Fig. S1). While some days experience more solar radiation before or after noon, there was no pattern consistent with morning clouds and afternoon sun. We do not expect preferential melting on the southwest sides of moss balls during the latter portion of July and early portion of August, 2009. Identical analysis using data from a boreal forest weather station site 20 km SE of our study site (RAWS site: May Creek, AK) revealed a very similar pattern of solar radiation to the Gates Glacier site, and the same lack of asymmetry in daily solar radiation timing. Thus, with the available data, we cannot explain the direction of moss ball motion.

### Persistence

We initially tagged 30 glacier moss balls in 2009. We subsequently recaptured 18 moss balls each in 2010, 2011, and 2012 (although this was not the same 18 moss balls each year). Recapture rates for individual moss balls were highly variable with some never seen again after the first year (*n* = 8) and others detected every year (*n* = 13). The best-fit survival model included differing apparent survival (□) among years, but with constant detection probability (*p*; Model 2; Table 1). This model received 58% of AICc weight, compared to 26% for the null model (Model 1), and less than 10% for the other models (Models 3 & 4; Table 1). The average annual rate of apparent survival, □, based on the null model, was 0.86 [95% confidence interval (CI) = 0.75-0.93], and the average detection rate was 0.84 (95% CI = 0.70-0.92). When parameterized by year, the annual apparent survival rate ranged from 0.74 in 2009-2010 to 1.0 in 2011-2012 with a particularly large 95% CI for 2010-2011 (Table 2; Fig. 4).

**Table 1.** Apparent survival models for glacier moss balls tested in this study with their corresponding Akaike’s Information Criterion scores that have been adjusted for small sample sizes (AICc). Relative AICc scores (ΔAICc) model weight are also given. Lower ΔAICc and higher model weight indicate greater support for a given model. Model components: probability of detection (*p*), apparent survival (ϕ).

| Model | Description | AICc | ΔAICc | Weight |
| --- | --- | --- | --- | --- |
| 1 | Null; No year effect on $p$ or $\phi$ | 107.09 | 1.56 | 0.26 |
| 2 | Year effect on $\phi$ | 105.53 | 0 | 0.58 |
| 3 | Year effect on $p$ | 108.92 | 3.39 | 0.10 |
| 4 | Year effect on both $p$ and $\phi$ | 110.25 | 4.72 | 0.05 |

**Table 2.** Estimates of the apparent survival (ϕ) and detection probability (*p*) of glacier moss balls for the two best-fit models. Parentheses after estimates indicate 95% confidence intervals.

| Model | Parameter | Estimate |
| --- | --- | --- |
| 1 | $p$ | 0.84 (0.70-0.92) |
| | $\phi$ | 0.86 (0.75-0.93) |
| 2 | $p$ | 0.82 (0.69-0.91) |
| | $\phi$ (2009-2010) | 0.74 (0.55-0.87) |
| | $\phi$ (2010-2011) | 0.98 (0.27-0.99) |
| | $\phi$ (2011-2012) | 1.0 (0.99-1) |

**Fig. 4.**
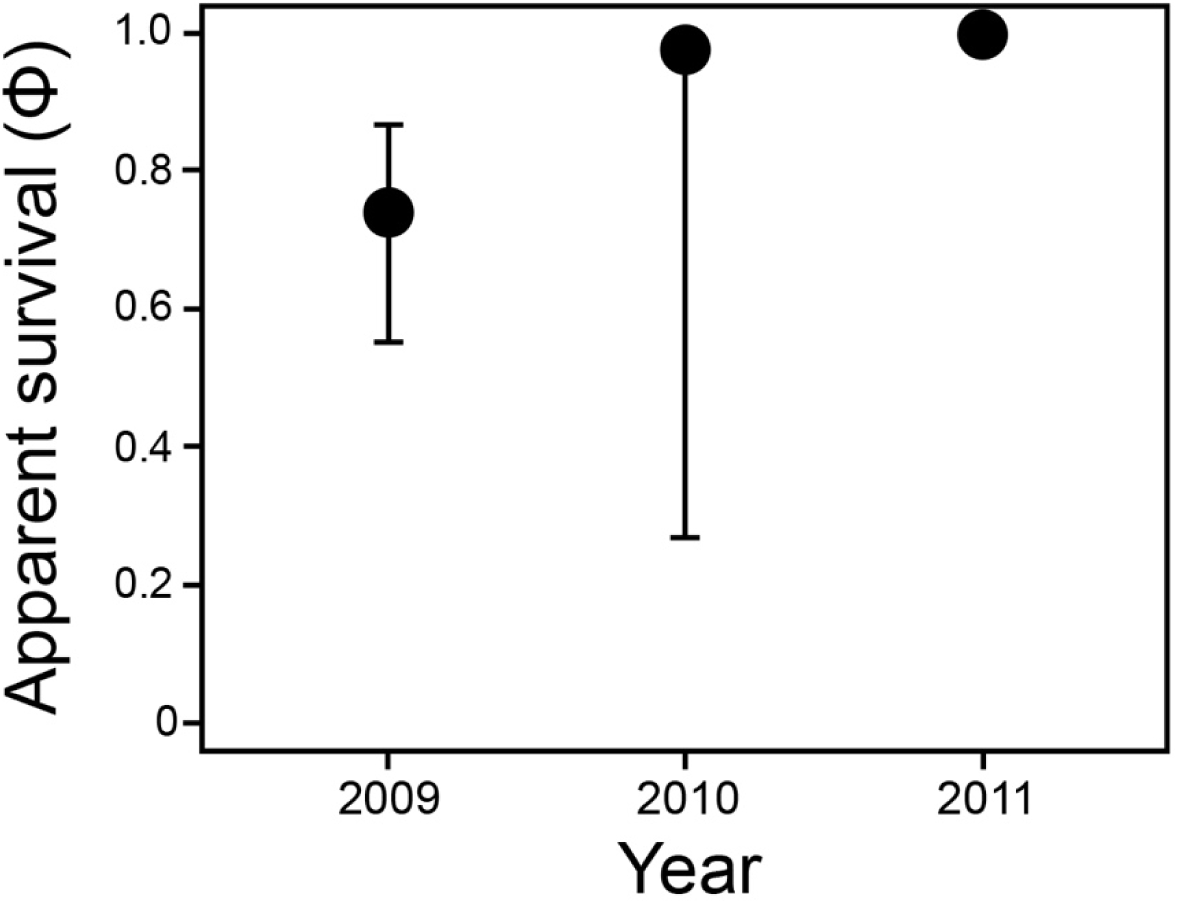
Estimates of apparent moss ball survival (□; dark circles) with 95% confidence intervals (thin dark lines) from model 2, the best-fit model, which included a year effect on □. Year-long, bracketed time intervals labeled on the x-axis are identified by their starting year. For instance, apparent survival for 2009-2010 is shown as 2009.

Our detection rate estimates may underestimate actual glacier moss ball survival for several reasons. First, at least four glacier moss balls lost their marking bracelet after the first year because we found the marking bracelet on the ice, separate from a moss ball. Second, another moss ball partially obscured its bracelet by growing to cover the beads, but we were able to detect a single bead and then delicately “excavate” the bracelet. Since we did not destructively search glacier moss balls that did not have an obvious bracelet, it is possible that additional instances of lost marking bracelets or growth to cover beads may have impacted our detection. Third, between 2009 and 2010, two tagged moss balls fell inside of a shallow crevasse within the study area. The two crevasse-bound glacier moss balls persisted, and likely continued to photosynthesize and grow to some capacity for the remainder of the study. We continued to check crevasses in the study area carefully, but some moss balls could have fallen into deeper crevasses, or into shallow crevasses in a way that obscured their markings, and therefore persisted without detection.

Our estimate of average life expectancy for a mature moss ball varied depending on whether the lowest overall or mean annual survival rate were used. If using the lowest annual survival rate (0.74), average life expectancy was 3.3 years (95% CI = 1.67-7.18). However, we expect this life expectancy to be biased low to some extent, because we were only able to estimate apparent survival (e.g., some insecure tags fell off moss balls that likely still persisted). If using the mean annual apparent survival rate across the entire study (0.86), average life expectancy rose to 6.63 years (95% CI = 3.48-13.78). This estimate may be biased high because we did not tag any new moss balls in years 2 and 3 (2010 and 2011), but simply re-captured existing (and therefore high survival probability) glacier moss balls. When thinking of lifespan, it is relevant to note that we also observed a glacier moss ball split roughly in half during the course of the study along its intermediate axis. The moss ball had become elongated and essentially pulled apart. This mechanism may contribute to keeping glacier moss balls ovular and represent a mode of moss ball genesis.

## Discussion

Glacier moss balls are intriguing components of glacier ecosystems that integrate physical (e.g., debris cover) and ecological (e.g., invertebrate colonization) factors into a unique habitat type. Previous research has revealed a great deal about glacier moss ball biology (e.g., their invertebrate colonizers, Coulson and Midgley 2012) yet their movement and longevity has remained unexplored. It has been speculated that glacier moss ball movement patterns likely follow the general downward slope of the glacier (Porter et al. 2008) and that they represent an ephemeral habitat type on glaciers, a factor that may limit colonization by specific invertebrate taxa (e.g., a lack of spiders; Coulson and Midgley 2012). Our results did not align with these predictions of movement and persistence.

### Movement

Even on the gently-sloped Root Glacier, glacier moss balls move relatively quickly (∼2.5 cm d^-1^) in similar directions and at similar speeds. Herd-like moss ball movements did not, however, follow the downward slope of the glacier, the dominant wind direction, nor the dominant direction of incoming solar radiation (Figs. 3, S1). Thus, we are left with a puzzling question: why do the azimuths of glacier moss balls appear to shift simultaneously throughout the summer season, resulting in the moss ball “herd” synchronously changing directions (Fig. 3a)? Moss balls began the season moving generally south and slowly transitioned towards the west. Given their movement independence from the dominant wind direction and downhill direction of the glacier, we speculated that shifts in patterns of solar radiation drive this pattern. Perhaps the weather transitioned from clear mid-day skies during late June and early July (associated with the most rapid motion and southerly azimuths), to a different weather pattern in late July of morning clouds and afternoon sun. Such a change could drive enhanced ablation on the west sides of moss balls, and therefore preferential westward movement. However, we found no evidence for diurnal solar radiation asymmetry during the study period (Fig. S1).

The relative contributions of downslope gravity versus another factor (e.g., solar radiation) almost certainly depend on glacier steepness. Porter et al. (2008) posited a considerable effect of gravity on glacier moss ball movement for a relatively steep (9.6°) Icelandic glacier which contrasts with our much flatter Root Glacier study area (∼3°). Still, regardless of steepness, differential melt patterns create pedestals that moss balls rest upon and, eventually, enough ice melts below the moss ball causing it to fall and flip (Porter et al. 2008). Assuming glacier moss balls are, on average, ∼10 cm in their intermediate axis, and their only means of movement is melt-induced flipping driven by pedestal emergence at the rate of 6-9 cm d^-1^, their rates of movement would imply each glacier moss ball flips every ∼2-4 days. However, we cannot rule out alternative modes of glacier moss ball movement. Many glacier moss balls have one side that is flattened and commonly faces down, while a more rounded, vegetated side faces skyward (Shacklette 1966). Given this orientation, an alternative scenario is that glacier moss balls also move by basal sliding over the wet glacier surface below.

### Persistence

Glacier moss balls persist across multiple years as stable ecological units. On average, 86% of the mature glacier moss balls included in this study survived annually which translates to a lifespan of more than 6 years. Thus, with high rates of survival across multiple years, and relatively high detection rates, we consider glacier moss balls to be long-lived, rather than ephemeral, glacier features. Unlike living individual organisms which can senesce as they age (e.g., Loison et al. 1999), moss ball survival rates are unlikely to decline with time in the traditional sense, nor should they exhibit density dependent survival (e.g., Festa-Bianchet et al. 2003). However, unlike traditional systems, factors that control disaggregation are likely the key process underlying moss ball longevity. The temporal stability of moss balls means they could exist for long enough to develop complex biotic communities (e.g., Coulson and Midgley 2012). However, the degree to which geographic location (e.g., distance to a glacier margin), and not persistence, influences invertebrate colonization remains to be tested.

The limited scope of our mark-recapture data collection precludes us from drawing conclusions about the inter-annual drivers of moss ball apparent survival. However, we can highlight factors that may influence it. First, it is possible that glacier moss balls moved more frequently out of the study area in one year versus others, perhaps due to exceptionally clear skies (and thus higher rates of glacier ablation). Second, we observed a number of fragmented moss balls. Fragmentation may be a normal part of moss ball growth trajectories, too frequent or intense freeze thaw cycles, or an as yet unknown factor. If glacier moss balls did survive within our study area, they had an 84% probability of being detected in a given year. This indicates that our bracelet and colored beads marking scheme was effective. However, for future studies, more robust marks should be considered (e.g., passive integrated transponder, PIT; Castro-Santos et al. 1996).

### Genesis, growth, and disaggregation

Our results allow us to add new speculation about patterns of glacier moss ball growth as well as additional evidence for previous hypotheses regarding their genesis and disaggregation (e.g., Heusser 1972; Perez 1991). In terms of growth, our documentation of glacier moss balls rolling over a fine-grained, wet, sedimentary substrate is consistent with growth through adherence of sediment to an existing moss ball. We observed “dirty” moss on some glacier moss balls in our study area. As the moss itself grows, this adhered sediment may become integrated within the fibrous material, increasing the size of the glacier moss ball. Field observation of moss growth over and around our identification bracelets indicates that several millimeters of growth can occur within years. However, the observation that most bracelets were not engulfed by sediment accumulation and moss growth during our four-year study period suggests either generally slow growth or an upper limit on moss ball size.

Understanding year-to-year moss ball growth, however, does not explain moss ball genesis, nor disaggregation. It is well-established that fibrous moss provides the skeletal structure that allows moss balls to be cohesive, ovoid structures. A source of moss spores is therefore essential to moss ball genesis (in our study, putatively, the Donoho nunatak). The question, then, is how glacier moss balls begin to grow in the first place, and on what substrate. Eythórsson (1951) suggested that a “stone kernel” at their centers is key. However, later investigations (e.g., Shacklette 1966; Coulson and Midgley 2012) found mixed results that largely reflected a consensus that there is no general rule about rock cores at the center of glacier moss balls. Our exploratory testing of moss balls also indicated that some, but not all, moss balls contained a ∼1-cm gravel “kernel” at their centers. Potentially, these kernels, with adhered fine-grained sediment, provide a growth substrate for initially wind-deposited moss spores. In our study area, the co-occurrence of moss balls within an unusually extensive, fine-grained “plume” of sediment cover (Fig. 1b) aligns with a similar observation by Heusser (1972) for the Gilkey Glacier in southeastern Alaska, USA. The origin of this fine-grained sediment is unknown, but in satellite imagery (Fig. 1b), it appears to originate from the ice itself and may be a volcanic ash layer being carried down from the high, volcanic, Wrangell Mountain peaks.

We identified few glacier moss balls greater than ∼15 cm on their long axis. Generally, moss balls appear to rarely exceed ∼10 cm except for rare cases in Alaska where they have been reported up to 18 cm (Benninghoff 1955; Heusser 1972). Why glacier moss balls in Alaska appear to grow larger than elsewhere in the world remains an open question but, regardless of location, there appears to be some size limiting process within the moss ball lifecycle. Shacklette (1966) suggested that the tensile strength of moss stems may be key. Exceeding this tensile limit may occur when the moss ball major axis grows too great relative to the intermediate axis. For instance, when a moss ball becomes too elongated, subtle variations in ice surface topography may lead the two ends of a moss ball to move in different directions and tear in the middle. During our study, we observed a splitting of a long, linear moss ball. While this process applies an upper-limit to moss ball size it also circles back to inform questions regarding the presence of a rock kernel. If the upper size limit is reached and a moss ball splits, only one of the two remaining moss balls involved in this “cloning” process will retain the gravel kernel. This may explain why a number of moss balls do not appear to have any coarse-grained rock at their cores. However, it is worth noting that in the case of Coulson and Midgley (2012), none of the moss balls in the study had a rock core. Therefore, glacier moss balls can almost certainly form without a “seed” rock.

## Conclusions

In this study, we extended previous research on glacier moss balls to quantify their movement and persistence on an Alaskan glacier. We showed that glacier moss balls move relatively quickly, at a rate of centimeters per day, in herd-like fashion. However, we could not explain the direction of moss ball movement by only considering the physical surface of the glacier (i.e., the downslope direction), the intensity of glacier ice ablation, and patterns of solar radiation. Thus, it appears a still unknown external force influences glacier moss ball movement on the Root Glacier. We also showed that mature moss balls are long-lived, with an average life expectancy of more than 6 years. The potential for glacier moss balls to act as relatively stable, long-term ecological units highlight their potential to act as key biotic habitat. Coulson and Midgley (2012) previously described invertebrate colonization of glacier moss balls and suggested that a lack of Enchytraeidae and Aranea may be the result of the ephemerality of moss balls in glacier habitats. Our results contrast this idea. Instead, we postulate that selective invertebrate colonization of glacier moss balls depends instead on their locations and frequent movements or, as Coulson and Midgley (2012) noted, the variable dispersal capacities of colonizers. Given the importance of microbial diversity to carbon cycling (Anesio et al. 2009), ecosystem function (Anesio et al. 2017; Hotaling et al. 2017a,b), and even albedo (Ganey et al. 2017), future efforts to understand the microbial ecology of glacier moss balls will further illuminate their ecological role in glacier ecosystems. Like cryoconite, the granular, darkly pigmented dust on glacier surfaces that drive hotspots of microbial activity (Cook et al. 2016), glacier moss balls may have similar value at the ecosystem scale.

## Acknowledgements

S.H. was supported by NSF award #OPP-1906015. We thank the Wrangell Mountains Center for logistical support and assisting with field measurements, and Dr. Billy Armstrong for providing the orthoimage of the study area.

## Compliance with Ethical Standards

The authors declare no conflicts of interest.

## References

Akaike H (1998) Information theory and an extension of the maximum likelihood principle. In: Selected papers of Hirotugu Akaike. Springer, New York, pp 199–213

Anesio AM, Hodson AJ, Fritz A, Psenner R, Sattler B (2009) High microbial activity on glaciers: importance to the global carbon cycle. Glob Change Biol 15:955–960

Anesio AM, Laybourn-Parry J (2012) Glaciers and ice sheets as a biome. Trends Ecol Evol 27:219–225

Anesio AM, Lutz S, Chrismas NA, Benning LG (2017) The microbiome of glaciers and ice sheets. NPJ Biofilms Microbiomes 3:1–11

Armstrong WH, Anderson RS, Allen J, Rajaram H (2016) Modeling the WorldView-derived seasonal velocity evolution of Kennicott Glacier, Alaska. J Glaciol 234:763–777

Belkina OA, Vilnet AA (2015) Some aspects of the moss population development on the Svalbard glaciers. Czech Polar Reports 5:160–175

Benninghoff WS (1955) Jökla-mýs. J Glaciol 2:514–515

Castro-Santos T, Haro A, Walk S (1996) A passive integrated transponder (PIT) tag system for monitoring fishways. Fish Res 28:253–261

Cook J, Edwards A, Takeuchi N, Irvine-Fynn T (2016) Cryoconite: the dark biological secret of the cryosphere. Prog Phys Geog 40:66–111

Coulson S, Midgley N (2012) The role of glacier mice in the invertebrate colonisation of glacial surfaces: the moss balls of the Falljökull, Iceland. Polar Biol 35:1651–1658

Deevey Jr ES (1947) Life tables for natural populations of animals. Q Rev Biol 22:283–314

Dial RJ, Becker M, Hope AG, Dial CR, Thomas J, Slobodenko KA, Golden TS, Shain DH (2016) The role of temperature in the distribution of the glacier ice worm, Mesenchytraeus solifugus (Annelida: Oligochaeta: Enchytraeidae). Arct Antarc Alp Res 48:199–211

Eythórsson J (1951) Correspondence. Jökla-mys. J Glaciol 1:503

Festa-Bianchet M, Gaillard JM, Côté SD (2003) Variable age structure and apparent density dependence in survival of adult ungulates. J Anim Ecol 72:640–649

Ganey GQ, Loso MG, Burgess AB, Dial RJ (2017) The role of microbes in snowmelt and radiative forcing on an Alaskan icefield. Nat Geosci 10:754–759

Gardner AS, Moholdt G, Cogley JG, Wouters B, Arendt AA, Wahr J, Berthier E, Hock R, Pfeffer WT, Kaser G (2013) A reconciled estimate of glacier contributions to sea level rise: 2003 to 2009. Science 340:852–857

Heusser CJ (1972) Polsters of the moss *Drepanocladus berggrenii* on Gilkey Glacier, Alaska. Bul Torrey Bot Club 99:34–36

Hotaling S, Hood E, Hamilton TL (2017a) Microbial ecology of mountain glacier ecosystems: biodiversity, ecological connections and implications of a warming climate. Environ Microbiol 19:2935–2948

Hotaling S, Finn DS, Joseph Giersch J, Weisrock DW, Jacobsen D (2017b) Climate change and alpine stream biology: progress, challenges, and opportunities for the future. Biol Rev 92:2024–2045

Hotaling S, Shain DH, Lang SA, Bagley RK, Lusha M, Weisrock DW, Kelley JL (2019) Long-distance dispersal, ice sheet dynamics, and mountaintop isolation underlie the genetic structure of glacier ice worms. Proc R Soc B 286:20190983

Hotaling S, Wimberger PH, Kelley JL, Watts HE (2020) Macroinvertebrates on glaciers: a key resource for terrestrial food webs? Ecology 101:e02947

Hurvich CM, Tsai C-L (1989) Regression and time series model selection in small samples. Biometrika 76:297–307

Larsen C, Burgess E, Arendt A, O’neel S, Johnson A, Kienholz C (2015) Surface melt dominates Alaska glacier mass balance. Geophys Res Lett 42:5902–5908

Lebreton J-D, Burnham KP, Clobert J, Anderson DR (1992) Modeling survival and testing biological hypotheses using marked animals: a unified approach with case studies. Ecol Monograph 62:67–118

Loison A, Festa-Bianchet M, Gaillard J-M, Jorgenson JT, Jullien J-M (1999) Age-specific survival in five populations of ungulates: evidence of senescence. Ecology 80:2539–2554

Mann D, Edwards J, Gara R (1980) Diel activity patterns in snowfield foraging invertebrates on Mount Rainier, Washington. Arct Antarc Alp Res 12:359–368

Millar JS, Zammuto RM (1983) Life histories of mammals: an analysis of life tables. Ecology 64:631–635

Perez FL (1991) Ecology and morphology of globular mosses of Grimmia longirostris in the Paramo de Piedras Blancas, Venezuelan Andes. Arct Antarc Alp Res 23:133–148

Porter P, Evans A, Hodson A, Lowe A, Crabtree M (2008) Sediment–moss interactions on a temperate glacier: Falljökull, Iceland. Ann Glaci 48:25–31

Roe GH, Baker MB, Herla F (2017) Centennial glacier retreat as categorical evidence of regional climate change. Nat Geosci 10:95–99

Rosvold J (2016) Perennial ice and snow-covered land as important ecosystems for birds and mammals. Journal Biogeogr 43:3–12

Shacklette HT (1966) Unattached moss polsters on Amchitka Island, Alaska. The Bryologist 346-352

Stibal M, Bradley JA, Edwards A, Hotaling S, Zawierucha K, Rosvold J, Lutz S, Cameron KA, Mikucki JA, Kohler TJ, Šabacká M, Anesio AM (2020) Glacial ecosystems are essential to understanding biodiversity responses to glacier retreat. Nat Ecol Evol https://doi.org/10.1038/s41559-020-1163-0

Uetake J, Tanaka S, Hara K, Tanabe Y, Samyn D, Motoyama H, Imura S, Kohshima S (2014) Novel biogenic aggregation of moss gemmae on a disappearing African glacier. PloS One 9:e112510

Van der Walt S, Schönberger JL, Nunez-Iglesias J, Boulogne F, Warner JD, Yager N, Gouillart E, Yu T (2014) scikit-image: image processing in Python. PeerJ 2:e453

