## Supplementary Materials for "Rolling stones gather moss: Movement and longevity of moss balls on an Alaskan glacier"

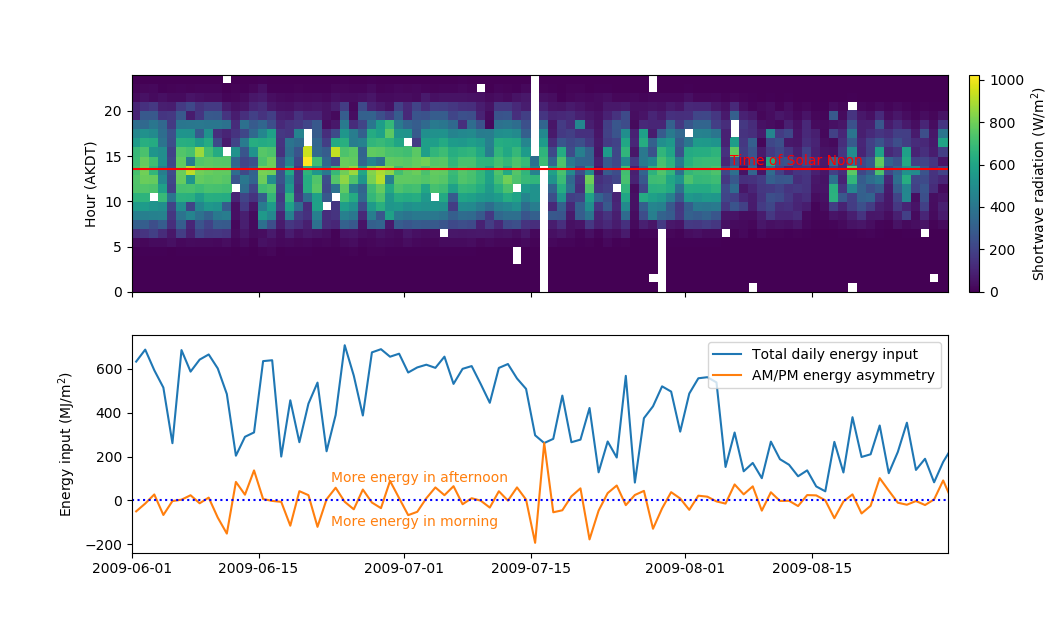

Fig. S1. Temporal variation in incoming solar radiation at Gates Glacier, Alaska, 15 km up-glacier of the moss ball study site. (Top panel) Shortwave radiation measured hourly as a function of hour and day, during three months including the measurement period for moss ball motion (18 June to 11 August 2009). Intervals of missing data are shown in white. Solar noon at the site, approximately 13:30 AKDT (21:30 UTC), is shown as a horizontal red line. (Bottom Panel) Total daily energy input and energy asymmetry over the same time interval illustrated in the top panel. In orange, days with more energy measured during the morning are shown as negative energy inputs and days with more energy measured after solar noon are shown with positive energy inputs. Missing data values are handled as zero values for the purpose of these integrated quantities.
